# Multi-omic characterization of the sow colostrum and milk microbiome and proteome

**DOI:** 10.1101/2025.11.05.686862

**Authors:** Devin B. Holman, Katherine E. Gzyl, Arun Kommadath, Pekka Määttänen

## Abstract

Sow colostrum and milk provide essential nutrients, immune protection, and one of the earliest microbial exposures for piglets. However, the microbial composition, functional potential, and host interactions of these mammary secretions remain poorly characterized. Here, we combined culturomics, metagenomics, and proteomics to comprehensively characterize the microbiome and proteome of sow colostrum and milk collected at farrowing and at 7 and 21 days postpartum. We recovered 132 bacterial isolates representing at least 42 species, including 15 putative novel taxa. These isolates included both potentially pathogenic species such as *Sarcina perfringens* and *Streptococcus suis* and potentially beneficial bacterial species like *Lactobacillus amylovorus* and *Lactiplantibacillus plantarum*. The microbial composition and functional potential shifted significantly as the milk matured, with *L. amylovorus*, *Limosilactobacillus reuteri*, and *Rothia* spp. among the most relatively abundant taxa. Several antimicrobial resistance genes, including *erm*(C), *tet(K)*, *tet*(M), *lnu(A)*, *poxtA*, and *fexB*, were identified on contigs encoding plasmid replicons in the isolates, indicating potential for horizontal gene transfer. Functional annotation of isolate genomes indicated broad carbohydrate-active enzyme (CAZyme) repertoires, including those conferring β-galactosidase activity and the capacity to metabolize milk oligosaccharides. The colostrum and milk proteome also shifted during lactation, reflecting declining immune-related proteins and increasing metabolic and structural proteins. Correlations between specific microbial taxa and host proteins, including *Rothia* spp. and immune proteins or glycoproteins, suggested potential host–microbe interactions during lactation. Together, these findings provide a multi-omic perspective on how mammary microbiome dynamics and host responses during lactation may influence neonatal microbial colonization and health.

## INTRODUCTION

Sow colostrum and milk serve as the primary sources of nutrition for piglets prior to weaning. Colostrum is the initial secretion from the mammary gland, typically produced within the first 24 h of farrowing, providing piglets with energy, as well as passive immunity in the form of immunoglobulins (IgA, IgG, and IgM) [1]. As mammary secretions transition to mature milk, the protein content decreases and the concentrations of lipids and lactose increase [2]. In addition to immunoglobulins, colostrum and milk contain a variety of bioactive proteins, including antimicrobial peptides such as lactoferrin, along with enzymes and signaling factors that can directly influence microbial colonization and immune system maturation [3]. Microorganisms are also present in colostrum and milk, and are among the earliest microbial exposures for piglets [4–7].

The origin of these colostrum- and milk-associated microbes in sows and other mammals remains a subject of debate. One proposed mechanism is oro/entero-mammary translocation, whereby microbes from the gastrointestinal tract or oral cavity are transported via immune cells to the mammary gland [8]. It is also possible that microbes on the skin and in the oral cavity of the offspring may enter the mammary gland (retrograde transfer) [8]. Regardless of their origin, colostrum and milk are important sources of early-life microbial exposure for piglets that shape the neonatal gut microbiome, modulate immune system development, and influence growth and health outcomes [9]. Therefore, a clearer understanding of the composition and functional potential of the sow colostrum and milk microbiome is needed to evaluate how these microbes may contribute to piglet health and dissemination of antimicrobial resistance.

Previous studies using 16S rRNA gene sequencing have identified members of the *Clostridium*, *Corynebacterium*, *Lactobacillus*, *Streptococcus*, and *Staphylococcus* genera as relatively abundant in sow colostrum and milk [10, 11]. However, these studies are limited by the inherent constraints of 16S rRNA gene sequencing with respect to both functional and taxonomic resolution. Higher-resolution metagenomic sequencing can mitigate some of these limitations by enabling species- and strain-level identification and functional gene profiling. Integrating microbiome and proteomic data can also provide insight into the host-microbe interactions in the mammary gland. In the present study, we used both metagenomic sequencing and culturomics to profile sow colostrum and milk at the species and strain levels, and integrated these data with proteomic analyses of the same samples. We hypothesized that sow colostrum and milk would contain potentially beneficial bacteria as well as other taxa that may serve as reservoirs of antimicrobial resistance genes (ARGs), and that the proteome would include host proteins associated with immune function and microbial colonization.

## METHODS

### Animals and experimental design

Pigs were cared for in accordance with the guidelines of the Canadian Council on Animal Care. All procedures and protocols involving animals were reviewed and approved by the Lacombe Research and Development Centre Animal Care Committee (animal use protocol number 201905). Oxytocin was administered to Landrace x Yorkshire sows (n = 14) inseminated with Duroc semen to ensure farrowing within the same 24-h period. Colostrum (day of farrowing) and milk samples (days 7 and 21 post-farrowing) were collected into sterile 150-mL screw-cap containers using sterile gloves, following thorough cleaning of the teat with 0.5% hydrogen peroxide (Prevail disinfectant wipes, Virox Technologies Inc., Oakville, ON, Canada). Samples were stored on ice and transported to the laboratory within 3 h of collection. The sows were housed in individual farrowing stalls and did not receive any antimicrobials during gestation or lactation.

### Bacterial isolation

Colostrum and milk samples were processed for culturing in a vinyl anaerobic chamber (Coy Laboratory Products, Grass Lake, MI, USA) supplied with a premixed gas of 10% H_2_, 10% _CO2,_ and 80% N_2_. Samples were diluted 10-fold in phosphate-buffered saline (PBS; Thermo Scientific, Mississauga, ON, Canada) containing 0.1 µM resazurin sodium salt (MilliporeSigma, Oakville, ON, Canada) and 8 mM sodium sulfite. From this dilution, 100 µL was spread onto agar plates with de Man, Rogosa, and Sharpe (Thermo Scientific), Wilkins-Chalgren Anaerobe (Thermo Scientific), Tryptone Soya (TS) (Thermo Scientific), or TS supplemented with 5% sterile rumen fluid (Bar Diamond, Parma, ID, USA). Plates were incubated anaerobically at 39°C for 48 h using an AnaeroGen system (Thermo Scientific), corresponding to the body temperature of sows. At least one colony was selected from each agar type based on morphology using a sterile toothpick and transferred to a screw-cap tube containing 350 µL of the corresponding broth with 15% glycerol. The tube caps were loosened and the tubes incubated anaerobically at 39°C for 48 h. Caps were then tightened and cultures stored at -80°C until reculturing.

### Bacterial isolate DNA extraction

Bacterial isolates were regrown on their original agar with the exception that TS with 5% rumen fluid was replaced with TS agar. Isolates were recultured under the same conditions used for the initial isolation. A single colony was then selected and inoculated into a Hungate tube (Chemglass Life Sciences, Vineland, NJ, USA) containing 10 mL of the corresponding broth for 16 h at 39°C. A DNeasy Blood & Tissue kit (Qiagen, Mississauga, ON, Canada) was used to extract genomic DNA from each culture according to the manufacturer’s protocol.

### Initial bacterial isolate screening

A total of 271 isolates were recovered and screened via Sanger sequencing of the near-full-length 16S rRNA gene. The 16S rRNA gene was amplified using the primers 8F (5′-AGA GTT TGA TCC TGG CTC AG-3′) and 1492R (5′-CGG TTA CCT TGT TAC GAC TT-3′) [12] together with a HotStarTaq Plus Master Mix kit (Qiagen). Briefly, in a total reaction volume of 20 µL, 0.2 µM of each primer was included along with 1X CoralLoad Concentrate, 2 ng of template DNA, and 1X HotStarTaq Plus Master Mix in molecular-grade water. The PCR conditions consisted of an initial 95°C heat activation step for 5 min, followed by 35 cycles of denaturation at 94°C for 1 min, annealing at 55°C for 1 min, and extension at 72°C for 2 min, with a final extension at 72°C for 10 min. The PCR amplicons (20 µL) were electrophoresed on a 0.8% agarose gel (Invitrogen, Waltham, MA, USA) for 45 min at 100 V. A single ∼1,500 bp band was excised from the gel and purified with a QIAquick gel extraction kit (Qiagen) as per the manufacturer’s instructions. Each purified PCR product was sent to Eurofins (Louisville, KY, USA) for Sanger sequencing with the 16S F standard primer option. The 16S rRNA gene sequences were then identified using BLASTn [13].

### Whole-genome sequencing of bacterial isolates

Based on the 16S rRNA gene classification, isolates representing lactic acid-producing species, potentially pathogenic species such as *Streptococcus suis*, potentially novel species, and at least one representative of each additional species identified, were selected for whole-genome sequencing. In addition to the isolates screened via Sanger sequencing of the 16S rRNA gene, 53 isolates were included without this screening step, resulting in a total of 132 isolates subjected to whole-genome sequencing. Genomic libraries were prepared using an Illumina DNA Prep Library kit as previously described [14], and sequenced on an Illumina MiSeq instrument (v3 reagent kit: 600 cycles; v2 reagent kit: 300 cycles).

Long-read sequencing was also conducted on isolates representing potentially novel bacterial species and on at least one strain from species within the genera *Lactiplantibacillus*, *Lactobacillus*, *Latilactobacillus*, *Limosilactobacillus*, and *Weissella*. For PacBio HiFi sequencing, DNA was extracted with the Nanobind CBB kit (Pacific Biosciences, Menlo Park, CA, USA) following the HMW DNA extraction protocol for gram-positive bacteria provided by the manufacturer. To enhance cell lysis efficiency, cell pellets were resuspended in 10 µL of PBS containing 10 mg/mL MetaPolyzyme (MilliporeSigma) and incubated for 30 min at 35°C, followed by a 90-min incubation with 10 mg/mL lysozyme in STET buffer (8% sucrose, 50 mM Tris-HCl, 50 mM EDTA, 5% Triton X-100). Genomic DNA from these isolates was purified using a DNeasy PowerClean Pro CleanUp kit (Qiagen) and sequencing libraries were prepared with the SMRTbell Prep kit 3.0 (Pacific Biosciences) following the manufacturer’s instructions.

Briefly, purified DNA (final volume: 65 µL) was sheared using a Megaruptor 3 instrument (Diagenode Inc., Denville, NJ, USA). Sequencing primer 3.2 was annealed and the Sequel II 3.2 polymerase was bound. Libraries underwent AMPure bead cleanup following the SMRT Link v. 11.0 calculator procedure before sequencing on a PacBio Sequel II instrument at a loading concentration of 90 pM, using the adaptive loading protocol with the Sequel II Sequencing kit 2.0, SMRT Cell 8M, and 15 h movies with a 2-h pre-extension time. For isolates sequenced on the MinION R10.4.1 flow cell (Oxford Nanopore, Oxford, UK), DNA extraction, library preparation, and sequencing were done as described previously [15].

### Bacterial genome assembly and analysis

Illumina raw reads were processed, quality-filtered, and assembled as described in Holman et al. [15] using the same parameters and software versions. Similarly, the processing and assembly of Nanopore reads were performed as previously detailed [15]. Processing and assembly of the PacBio reads were similar to those used for the Nanopore reads with the exception that the minimum read length was set to 1,000 bp in Filtlong v.0.2.1 (https://github.com/rrwick/Filtlong) and Medaka polishing was omitted in Trycycler v.0.5.4 [16]. Assemblies were identical before and after polishing, except for *Lactobacillus amylovorus* 11266D007BMRS-1, for which the short-read-polished assembly was retained. For all genome assemblies, taxonomy was assigned using the Genome Taxonomy Database toolkit (GTDB-Tk) v.2.4.1 and the GTDB release 10-R226 [17]. CheckM2 v.1.0.1 [18] was used to assess completeness and contamination, and QUAST v.5.2.0 [19] was used to determine assembly metrics. Antimicrobial resistance genes (ARGs) were identified with the resistance gene identifier (RGI) v.6.0.3 and the Comprehensive Antibiotic Resistance Database (CARD) v.4.0.0 [20]. Screening for plasmid replicons was done with PlasmidFinder v.2.1.6-1 [21]. The average nucleotide identity (ANI) between isolates was calculated using fastANI v.1.33 [22].

### Metagenomic DNA extraction, sequencing, and analysis

Details of DNA extraction from colostrum and milk samples, metagenomic library preparation, initial sequence processing are available in Holman et al. [15]. Kraken2 v.2.1.6 [23] and Bracken v.3.0.1 [24] with the GTDB release 10-R226 were used to classify the unassembled sequences. The Kraken2- and Bracken-formatted GTDB also included the pig (Sscrofa11.1) and human (GRCh38.p14) genomes. Antimicrobial resistance genes were identified with RGI v.6.0.3 and CARD v.4.0.0 with KMA. The metagenomes were assembled both individually for each sample and as a co-assembly using MEGAHIT v.1.2.9 [25]. Reads were then mapped back to each respective individual assembly as well as the co-assembly using Bowtie2 v.2.5.1 [26] and the contigs were binned into metagenome-assembled genomes (MAGs) with MetaBAT 2 v.2:2.15 [27]. The completeness and contamination of the MAGs were assessed with CheckM v.1.2.2 and those MAGs with >90% completeness and <5% contamination were retained. Redundant MAGs were removed with dRep v.3.4.3 [28] with primary and secondary clustering at 90% and 99 % ANI, respectively. A phylogenomic tree of the MAGs and isolate genomes was constructed using PhyloPhlAn v.3.1.68 [29] and visualized in iTOL v.6.9.1 [30].

The unassembled metagenomic reads were aligned to the Kyoto Encyclopedia of Genes and Genomes (KEGG) release 112.0 [31] prokaryotic protein database with DIAMOND v.2.1.8.162 [32]. Gene hits were assigned to KEGG orthology (KO) groups and KO abundances were normalized by sequencing depth. KO groups were then filtered with MinPath [33] to identify a parsimonious set of pathways, and the remaining KO groups were assigned to KEGG pathways. The relative abundances of MAGs and isolates in sow milk and colostrum samples were estimated using CoverM v.0.7.0 [34]. Reads from untreated and pre-weaned piglets from these sows were mapped to these MAGs and isolate genomes as described in Holman et al. [15]. Both the isolate genomes and MAGs were analyzed using run_dbcan v.5.2.1 with DIAMOND and PyHMMER v.0.11.0 [35] under default settings against the dbCAN3 database (release: 2025-09-13) [36] to annotate carbohydrate-active enzymes (CAZymes) and identify CAZyme gene clusters (CGCs) and their predicted polysaccharide substrates.

### Proteomics

The protein content of the colostrum and milk was assessed at the Proteomics Core Facility, Université de Montréal (Montreal, QC, Canada). An initial acetone precipitation was performed on 40 µL of each colostrum and milk sample. The samples were then centrifuged and the supernatant was removed. The pellet was reconstituted in 100 mM Tris-HCl (pH 8.2) containing 8 M urea, vortexed and centrifuged at 17,000 x g for 10 min. Supernatants were collected and protein concentrations were determined using the Bradford assay. The solution containing 100 µg of protein was diluted in 100 mM Tris-HCl with 1 M urea and 10 mM TCEP [Tris(2-carboxyethyl)phosphine hydrochloride; Thermo Scientific], and vortexed for 1 h at 37°C.

Chloroacetamide (Sigma-Aldrich) was then added to a final concentration of 40 mM for alkylation and the samples were vortexed for an additional hour at 37°C. One microgram of trypsin was then added, and digestion was performed for 8 h at 37°C. The samples were next desalted on ultra-microspin columns (The Nest Group Inc., Ipswich, MA, USA), dried and solubilized in 5% acetonitrile and 4% formic acid. The samples were then loaded onto a 1.5 µL pre-column (Optimize Technologies, Oregon City, OR, USA).

Peptides were separated on a homemade reversed-phase column (150 μm inner diameter x 200 mm) with a 56 min gradient from 10 to 30% acetonitrile and 0.2% formic acid and a 600 nL/min flow rate on an Easy-nLC 1200 system coupled to a Q Exactive HF Orbitrap LC-MS/MS System (Thermo Scientific). Each full mass spectrometry (MS) spectrum, acquired at a resolution of 120,000, was followed by acquisition of tandem mass spectra on the most abundant multiply charged precursor ions for 3 s. Tandem MS experiments were performed using higher-energy collision dissociation at a collision energy of 34%. The data were processed using PEAKS X Pro (Bioinformatics Solutions, Waterloo, ON) and the UniProt *S*. *scrofa* database. Mass tolerances on precursor and fragment ions were 10 ppm and 0.01 Da, respectively. Carbamidomethylation was used as a fixed modification. Variable post-translational modifications included acetylation, oxidation, deamidation, and phosphorylation. The data were visualized with Scaffold 5.0 using a protein threshold of 99%, with at least 2 peptides identified and a peptide-level false discovery rate (FDR) of 1%.

### Statistical analysis

All statistical analyses were conducted in R v.4.5.1. The composition of the microbiome (species and relative abundances), resistome (ARGs; counts per million reads [CPM]), and KEGG orthologs (KOs; CPM) were assessed by sampling day using Bray–Curtis dissimilarities and PERMANOVA, implemented in vegan v.2.7-1 [37]. MaAsLin3 v.1.1.0 [38] was used to identify differentially abundant ARGs, microbial species, and KEGG pathways between the three sampling times. The protein abundance table exported from Scaffold 5.0, which reports relative intensities, was analyzed after removing proteins deleted from UniProt and matching gene names to their corresponding UniProt entries. Principal component analysis (PCA) was carried out using PCAtools v.2.16.0 [39], after excluding the 10% least variable proteins. Scores for the first and second principal components, together with the top five contributing proteins for each component, were extracted and visualized with ggplot2 v.3.5.2 [40]. Gene names were dereplicated by retaining the first occurrence, and loading arrows were scaled to match the range of PCA scores.

Protein abundance values were normalized with cumulative sum scaling in the metagenomeSeq package v.1.46.0 [41] and then log_2_-transformed. Differential abundance analysis was performed with limma v.3.60.6 [42] by fitting a linear model across all samples with time included as a factor. A contrast matrix was constructed to compare day 0 vs. day 7, day 0 vs. day 21, and day 7 vs. day 21. Moderated t-statistics and *P*-values were obtained using empirical Bayes, and *P*-values were adjusted with the Benjamini-Hochberg (BH) procedure. Volcano plots were generated with thresholds of absolute log_2_ fold change greater than 1 and adjusted *P*-values below 0.05. For each comparison, the five most significant proteins were labeled with their UniProt gene names. Correlations between the microbiome (species) and proteome were evaluated in vegan via Procrustes analysis based on Bray–Curtis dissimilarity non-metric multidimensional scaling (NMDS) ordinations.w

Gene ontology (GO) biological process enrichment analysis was performed using the PANTHER GO Enrichment Analysis tool v. 19.0 [43] with default settings and the *S*. *scrofa* reference database. For each comparison, the input consisted of the differentially abundant proteins (UniProt entries) with a log₂ fold change greater than 0.5 and an FDR of less than 0.05. Enriched GO terms were then mapped to higher-level summary categories using the R packages ontologyIndex v.2.12, GO.db v.3.19.1, AnnotationDbi v.1.66.0, and the GOslim generic ontology database (v. July 22, 2025).

## RESULTS

### Bacteria isolated from sow colostrum and milk

The 132 isolates whose genomes were sequenced and classified belonged to at least 42 unique species within 3 phyla: Actinobacteriota, Bacillota, and Pseudomonadota (Fig. 1; Table S1). Fifteen isolates were identified as novel taxa or as species without cultured representatives. These belonged to the *Corynebacterium*, *Rothia*, *Staphylococcus*, and *Streptococcus* genera. The majority of isolates (n = 91) were classified as either *Staphylococcus* or *Streptococcus* spp., representing at least 19 different species.

**Figure 1.**
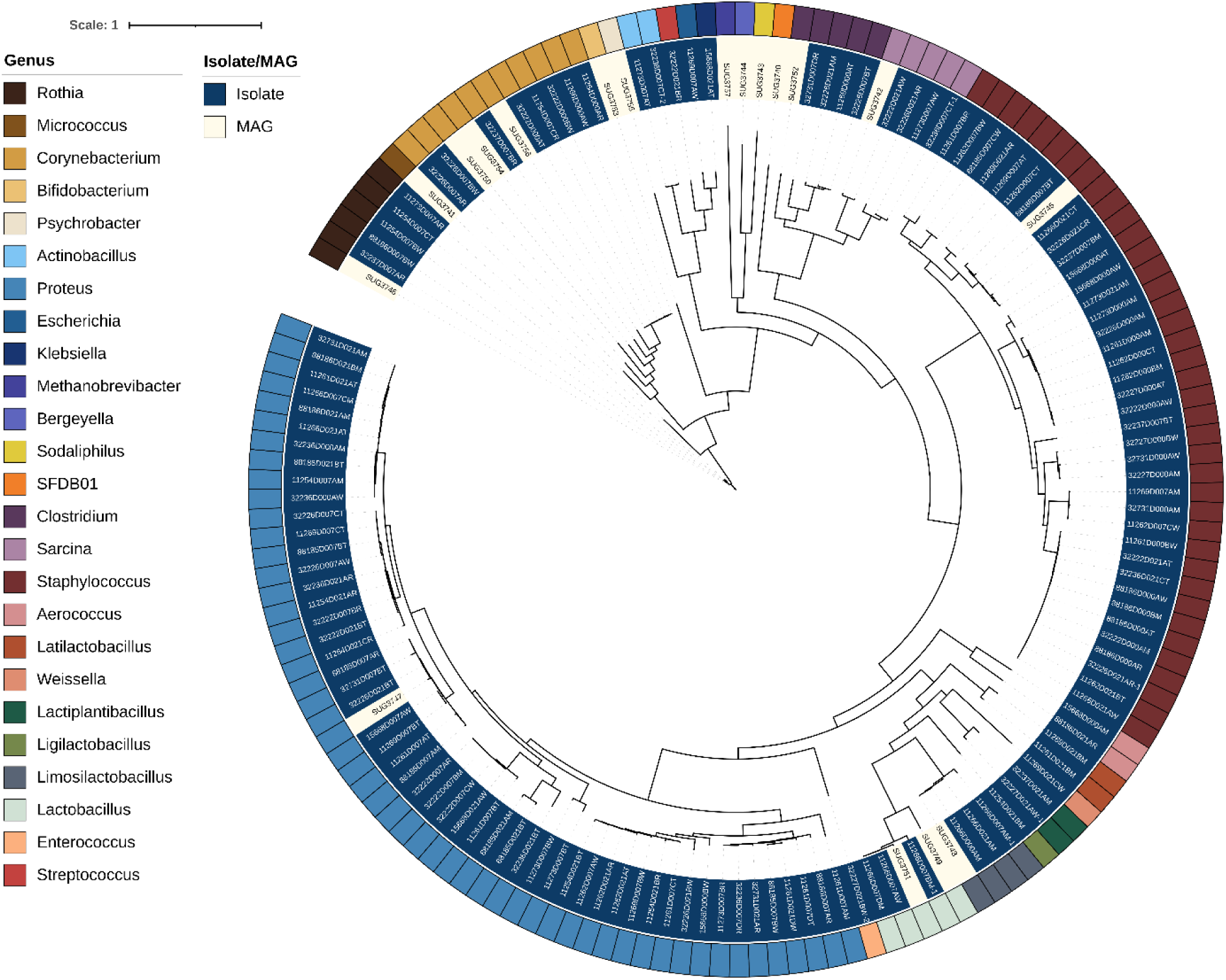
Maximum-likelihood phylogenetic tree of metagenome-assembled genomes (MAGs; inner ring, white) and isolate genomes (inner ring, blue). Genus-level classifications are indicated by outer ring colours.

One of the objectives of this study was to isolate *Lactobacillaceae* bacteria, as members of this family have been well established to have beneficial properties in various animal hosts, including pigs. Twelve *Lactobacillaceae* isolates were recovered and identified as *Lactiplantibacillus plantarum*, *L. amylovorus*, *Latilactobacillus curvatus*, *Limosilactobacillus reuteri*, or *Weissella paramesenteroides*. Potentially pathogenic or zoonotic strains were also recovered from colostrum and milk, including *Aerococcus suis*, *Escherichia coli*, *S*. *suis*, and *Sarcina* (*Clostridium*) *perfringens*. *In silico* serotyping identified the *E*. *coli* isolate as serotype O6:H7. Using a ≥ 99.99% ANI to define strains as recommended by Rodriguez-R et al. [44], there were potentially 12 unique *S*. *suis* strains represented among the isolate collection.

### Antimicrobial resistance genes in isolates

Apart from the multidrug efflux pump-encoding gene *sdrM,* which was found in all *Staphylococcus* isolates, the most frequently detected ARGs were *ant(6)-Ia*, *erm*(B), *lnu*(A), *lnu*(B), *lsa*(E), and *tet*(O) (> 20 isolates). With the exception of one isolate (11254D021BT), all *S*. *suis* isolates (n =13) carried at least one tetracycline resistance gene, including *tet*(40), *tet*(45), *tet*(L), *tet*(O), or *tet*(O/W/32/O). Notably, *L. curvatus* 11269D021BM carried two distinct ARG-encoding plasmids: a rep29 plasmid (p11269D021BM-1; 25,787 bp) harbouring *fexB* and *poxtA* and a rep28 plasmid (p11269D021BM-2; 10,359 bp) encoding *tet*(M) (Table S2). For plasmid p11269D021BM-1, the closest match in the NCBI nucleotide database was an *Enterococcus faecium* plasmid (99.71% identity, 88% coverage; accession CP072887.1) in an isolate recovered from surface water [45]. Plasmid p11269D021BM-2 aligned most closely with a *Lactiplantibacillus plantarum* plasmid (99.87% identity, 87% coverage; accession: AF440277.1) isolated from corn silage [46]. In addition to phenicols, the *poxtA* gene confers resistance to oxazolidinones [47], which include linezolid, an antibiotic of last resort used to treat serious infections caused by gram-positive bacteria such as methicillin-resistant *Staphylococcus aureus* (MRSA) and vancomycin-resistant enterococci (VRE) [48]. *Weissella paramesenteroides* 11269D021CW also carried a plasmid with *tet*(M), albeit larger (16,510 bp) and without a known replicon type.

Other ARGs located on the same contigs as plasmid replicon genes included *tet*(K) (rep7a) in several *Staphylococcus epidermidis* and *Staphylococcus simulans* isolates, and *tet*(M) (repUS43) in *Aerococcus suis*, *Staphylococcus hyicus*, and *Staphylococcus rostri* genomes (Table S2). The *lnu*(A) gene conferring resistance to lincosamides was associated with three different replicon types depending on the species and strain: rep13 (*S. simulans*), rep33 (*Streptococcus orisratti*, *Streptococcus thoraltensis*), and repUS54 (*Enterococcus dispar* 32227D021BWCA-2 and *L*. *curvatus* 11261D021BM). The aminoglycoside resistance gene *ant(4’)-Ia* was co-located with rep22 in *S*. *epidermidis* 11262D021BTSA.

### Carbohydrate-active enzymes in isolates

The isolate genomes were analyzed for CAZymes, CGCs, and their predicted polysaccharide substrates to examine their distribution and provide insight into the functional potential of the isolates. CAZymes catalyze the breakdown, modification, and synthesis of carbohydrates, while CGCs comprise physically linked genes encoding at least one CAZyme, a transporter, a transcriptional regulator, and a signal transduction protein involved in carbohydrate metabolism [49]. Glycoside hydrolases (GHs), which hydrolyze glycosidic bonds, are the largest class of CAZymes. The isolates with the greatest number of unique GH families were those classified as *S*. *perfringens, Clostridium nitritogenes, Klebsiella pneumoniae, E. coli,* and *S. suis* (Table S1). The most frequently encoded GHs were GH13 (98.5% of 132 genomes), GH32 (93.9%), GH1 (88.6%), GH73 (87.9%), and GH23 (84.8%). Another CAZyme class, the glycosyltransferases (GTs), which are involved in glycan synthesis, was also widespread; GT2, GT4, GT28, GT51, and GT119 were identified in all isolate genomes.

A total of 32 polysaccharides were predicted to serve as substrates for the detected CGCs. Starch, human milk oligosaccharides, fructan, trehalose, galactomannan, and raffinose were among the most common substrates, each associated with CGCs in at least 60 genomes. Nearly all genomes (93.9%) encoded at least one GH family (GH1, GH2, GH35, or GH42) with β-galactosidase activity, conferring the ability to hydrolyze lactose, the most abundant carbohydrate in sow colostrum and milk [3]. *Corynebacterium* sp028726465, *Corynebacterium* sp024580955, and *Rothia* sp034179845 were the only isolates lacking all of these GH families. In addition to GH2 and GH42, other GH families involved in the breakdown of milk oligosaccharides include GH20 (β-hexosaminidases), GH29 and GH95 (fucosidases), GH33 (sialidases), GH112 (galacto-N-biose/lacto-N-biose phosphorylases), and GH136 (lacto-N-biosidases) [50]. With the exception of GH136, the *S*. *perfringens* isolates encoded CAZymes from all of these families, indicating their ability to metabolize milk oligosaccharides.

### Sow colostrum and milk metagenomes

As reported previously, the vast majority of sequences in the colostrum and milk metagenomes originated from the host (97.1 ± 1.2%, SEM) [15]. The average number of reads per sample after quality-filtering and host sequence removal was 2,018,105 ± 896,045 (data not shown). Metagenomic reads from the colostrum and milk samples, as well as from feces collected from the piglets, were aligned to the isolate genomes to assess their relative abundance. For the colostrum, the *L. amylovorus* and *L. reuteri* genomes as well as two *Rothia* spp. genomes (88186D007BWCA and 32237D007ATSR) were the only isolates with a relative abundance greater than 0.1% (Supplementary Table S1). In the milk collected on days 7 and 21, all unidentified *Rothia* spp. isolates were relatively abundant (> 0.1%), along with isolates classified as *Aerococcus urinaeequi*, *L*. *amylovorus*, *L*. *reuteri*, *Sarcina ventriculi*, and *Staphylococcus hyicus*. The *E*. *coli* isolate was by far the most abundant in the piglet fecal microbiome (11.7 ± 0.8%) followed by *S*. *perfringens* and the *L*. *amylovorus* isolates (≥ 1.0%).

### Colostrum and milk microbiome composition, resistome, and functional potential

When the colostrum and milk metagenomic reads were profiled against the GTDB, the most relatively abundant species across all samples were *Methanocatella (*NCBI: *Methanobrevibacter)* sp900769095, *L*. *amylovorus*, *Rothia* sp034090985, *Rothia* sp034179845, and *S. suis* (Table S3). It is important to note that 24.1% ± 3.9% of the reads were assigned to the pig genome despite a host read removal step during pre-processing (data not shown). This highlights the importance of including host genomes in the Kraken2 database for taxonomic classification of low-biomass samples and is consistent with the findings of Gihawi et al. [51].

The microbial species composition of the colostrum and milk microbiomes differed significantly across all sampling time points (PERMANOVA: *R²* = 0.21, *P* < 0.001; Fig. 2a). The most pronounced shifts were observed between day 0 (colostrum) and day 7 (*R²* = 0.19, *P* < 0.001), as well as between day 0 and day 21 (*R²* = 0.18, *P* < 0.001). Significant differences were also detected between day 7 and day 21 (*R²* = 0.11, *P* < 0.001). A total of 23 species were identified as differentially abundant between day 0 and day 7, while 21 species differed between day 0 and day 21 (FDR < 0.05; Table S4). Of these, 13 species were consistently differentially abundant in both the day 0 vs. day 7 and day 0 vs. day 21 comparisons. The majority of these species were more abundant in the day 0 colostrum samples. Species enriched in the colostrum samples included *Aeromonas salmonicida*, *Lelliottia chinensis*, *Serratia proteamaculans*, and *Streptococcus faecavium*, whereas *Romboutsia timonensis*, *Rothia* sp034090985, *S. suis* (GTDB species cluster V), and *Turicibacter bilis* were more abundant in the day 7 and 21 milk samples. Additionally, 17 bacterial species were differentially abundant between the day 7 and 21 samples.

**Figure 2.**
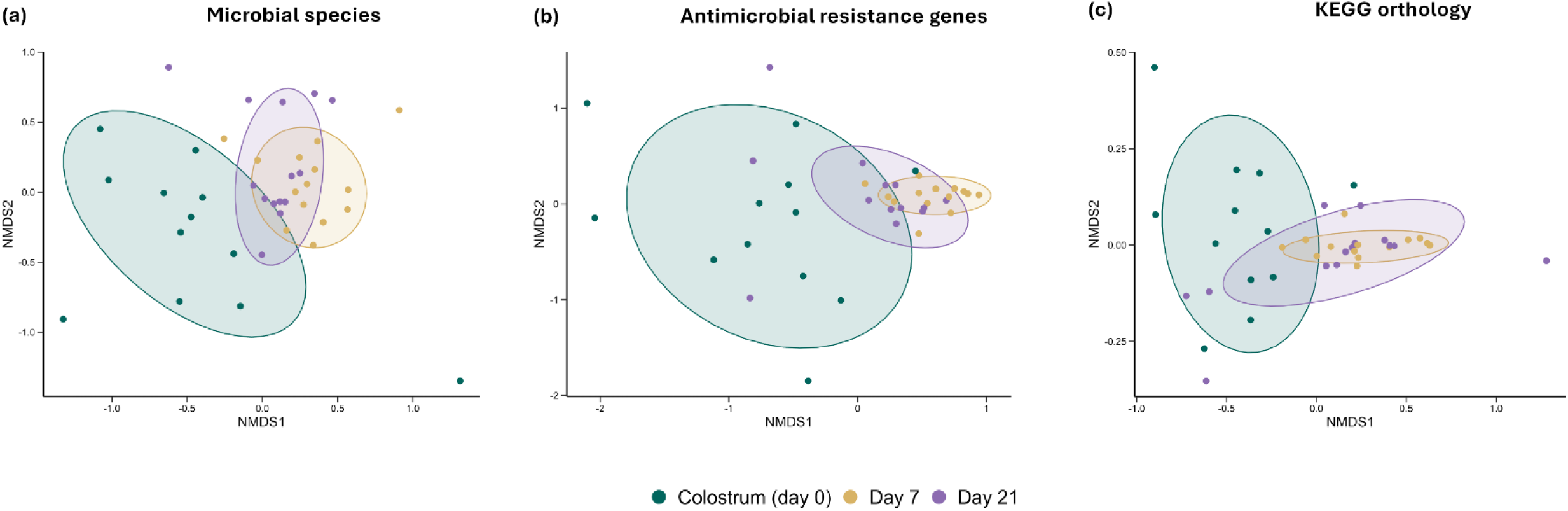
Non-metric multidimensional scaling (NMDS) ordinations of day 0 (colostrum), day 7, and day 21 samples based on Bray– Curtis dissimilarities of **(a)** archaeal and bacterial species relative abundances, **(b)** antimicrobial resistance gene abundances (counts per million reads, CPM), and **(c)** KEGG ortholog (KO) relative abundances (CPM). Points represent individual samples and ellipses denote 80% confidence intervals around group centroids.

Distinct shifts in resistome composition were also observed as milk matured (PERMANOVA: *R²* = 0.17, *P* < 0.001; Fig. 2b), with the most substantial changes between day 0 and day 7 (*R²* = 0.18, *P* < 0.001) and between day 0 and day 21 (*R²* = 0.12, *P* < 0.001). Despite these significant differences in overall composition, only a limited number of individual ARGs were differentially abundant between day 0 and days 7 and 21. Specifically, *lnu*(C) and *lsa*(E) were enriched in day 7 samples, while *tet*(40) was enriched in day 21 samples compared to day 0 (FDR < 0.05). The most relatively abundant ARGs across all samples were those conferring resistance to the tetracyclines such as *tet*(W), *tet*(Q), *tet*(K), and *tet*(O), and to the macrolides-lincosamides-streptogramin B class, including *lnu*(A) and *lnu*(C) (Table S5).

Similar to the species and ARG composition, the functional content (KO groups) of the milk microbiomes differed significantly between day 7 (*R^2^* = 0.18, *P* < 0.001) and day 0, as well as between day 21 and day 0 (*R^2^* = 0.11, *P* < 0.001) (Fig. 2c, NMDS). The KO groups were also assigned to KEGG functional pathways and compared between sampling times. There were 37 functional pathways enriched in the day 7 vs. day 0 samples and 32 in the day 21 vs. day 0 samples (FDR < 0.05). Notably, no functional pathways were more relatively abundant in the day 0 (colostrum) samples compared to the other two sampling periods, nor were any differentially abundant between the day 7 and day 21 samples (FDR > 0.05; Table S6). Pathways related to carbohydrate metabolism (e.g., phosphotransferase system, starch and sucrose metabolism, galactose metabolism), lipid metabolism (glycerophospholipid metabolism, biosynthesis of unsaturated fatty acids), and energy production (oxidative phosphorylation, nitrogen metabolism, porphyrin metabolism) were among those significantly more abundant in the microbiomes of day 7 and 21 milk.

### Metagenome-assembled genomes

The metagenomic reads were assembled and binned into 18 high-quality MAGs (≥ 90% completeness; ≤ 5% contamination) (Fig. 1; Table S7). Six of these MAGs corresponded to species also recovered in the isolate collection, including *L*. *amylovorus*, *L*. *reuteri*, *S*. *ventriculi*, *Rothia* sp034179845, *S*. *chromogenes,* and *S*. *hyovaginalis*. In the phylogenetic tree, all of these MAGs clustered with at least one isolate genome. For example, the *S*. *ventriculi* MAG (SUG3742) grouped with the S. *ventriculi* isolate (32222D021AW), although the *lnu*(P) and *tet*(O) genes identified in the isolate were not binned into the MAG (Fig. 1; Table S7). Likewise, *L*. *amylovorus* isolates 11266D007AW and 11266D007DM and MAG SUG3751 were highly similar, as were *S*. *hyovaginalis* isolates 32226D021BT, 32731D007BT, 88185D007AR, and MAG SUG3747.

Species recovered only as MAGs included *Bergeyella zoohelcum*, *Bifidobacterium mongoliense*, *Psychrobacter sanguinis,* and the archaeal placeholder *Methanocatella* sp900769095. The MAGs classified as *L*. *amylovorus*, *Rothia* sp034179845, and *L*. *reuteri* were most relatively abundant in both the colostrum and milk samples (Table S7). Within piglet fecal microbiomes, the *L*. *amylovorus, Lactobacillus* sp910589675*, L*. *reuteri*, and *Sodaliphilus* sp004557565 MAGs were most relatively abundant (> 0.1%). Although many ARGs detected in isolates were not binned into MAGs, *ant(6)-Ia, aph(3’)-Ia,* and *tet*(W/N/W) were detected in MAGs classified as *L*. *amylovorus, P. sanguinis,* and *L. reuteri*, respectively.

### Sow colostrum and milk proteomes

The proteomes of the colostrum and milk were analyzed to assess changes in protein composition during lactation and their potential relationship with the microbiome. Caseins and whey proteins such as albumin (ALB), α-lactalbumin (LALBA), β-lactoglobulin, and whey acidic protein, along with immune-related proteins, were relatively abundant across colostrum and milk samples (Table S8). As with the microbiomes, the proteomes of the colostrum and milk samples also differed significantly by time (Fig. 3a). Of the 3,680 proteins quantified, colostrum (day 0) and mature milk (day 21) exhibited the greatest divergence, with 681 proteins significantly more abundant in the colostrum and 523 proteins significantly more abundant in the mature milk (FDR < 0.05) (Fig. 3b; Table S9). Similarly, the transition from colostrum to early milk (day 0 vs. day 7) was marked by substantial differences, with 569 proteins significantly more abundant in day 0 samples and 469 in day 7 samples (FDR < 0.05). In contrast, relatively few differences were detected between early and mature milk (day 7 vs. day 21), with only 59 proteins significantly more abundant in the day 7 milk samples and 63 in the day 21 samples (FDR < 0.05).

**Figure 3.**
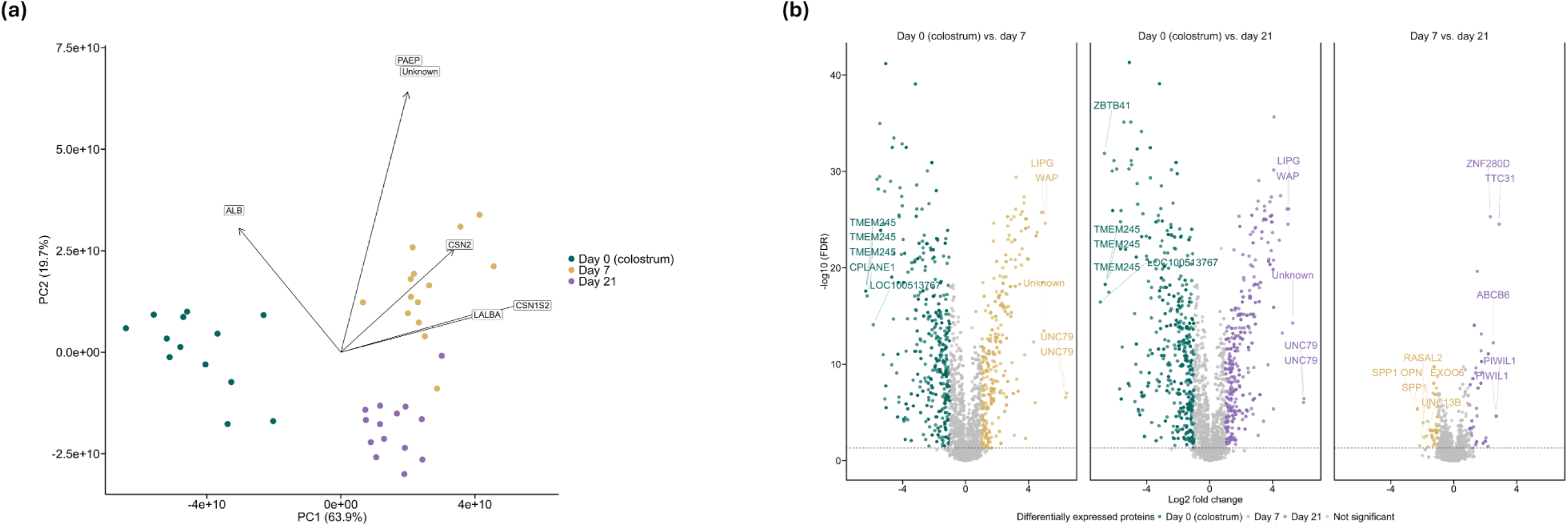
(a) Principal component analysis (PCA) biplot of colostrum (day 0) (n = 13), day 7 (n = 14), and day 21 (n = 14) samples based on proteomic profiles. Arrows indicate protein loadings that contribute most strongly to separation along the first two principal components. **(b)** Volcano plots of differentially expressed proteins for pairwise comparisons (colostrum vs. day 7, colostrum vs. day 21, and day 7 vs. day 21). Coloured points denote proteins enriched in each group, gray points indicate non-significant changes, and selected significant proteins are labeled. The horizontal dashed line marks the false discovery rate (FDR) threshold for significance.

Several of the whey proteins were among those differentially abundant between colostrum and mature milk (day 7 and 21) samples, with ALB, serotransferrin (TF), and haptoglobin (HP) enriched in the colostrum and LALBA and lactoperoxidase (LPO) enriched in the day 7 and 21 samples (Table S9). Among the proteins most strongly linked with colostrum samples were colostrum trypsin inhibitor-like protein, ciliogenesis and planar polarity effector complex subunit 1 (CPLANE1), and several zinc finger proteins. The differentially abundant proteins were assigned to GO terms for functional annotation. Among the most enriched GO terms in colostrum were those related to immune system processes and response, as well as wound healing and cell differentiation (Fig. 4; Table S10). Many of the GO terms enriched among the day 7 and 21 samples compared to day 0 were those associated with cytoskeleton organization.

**Figure 4.**
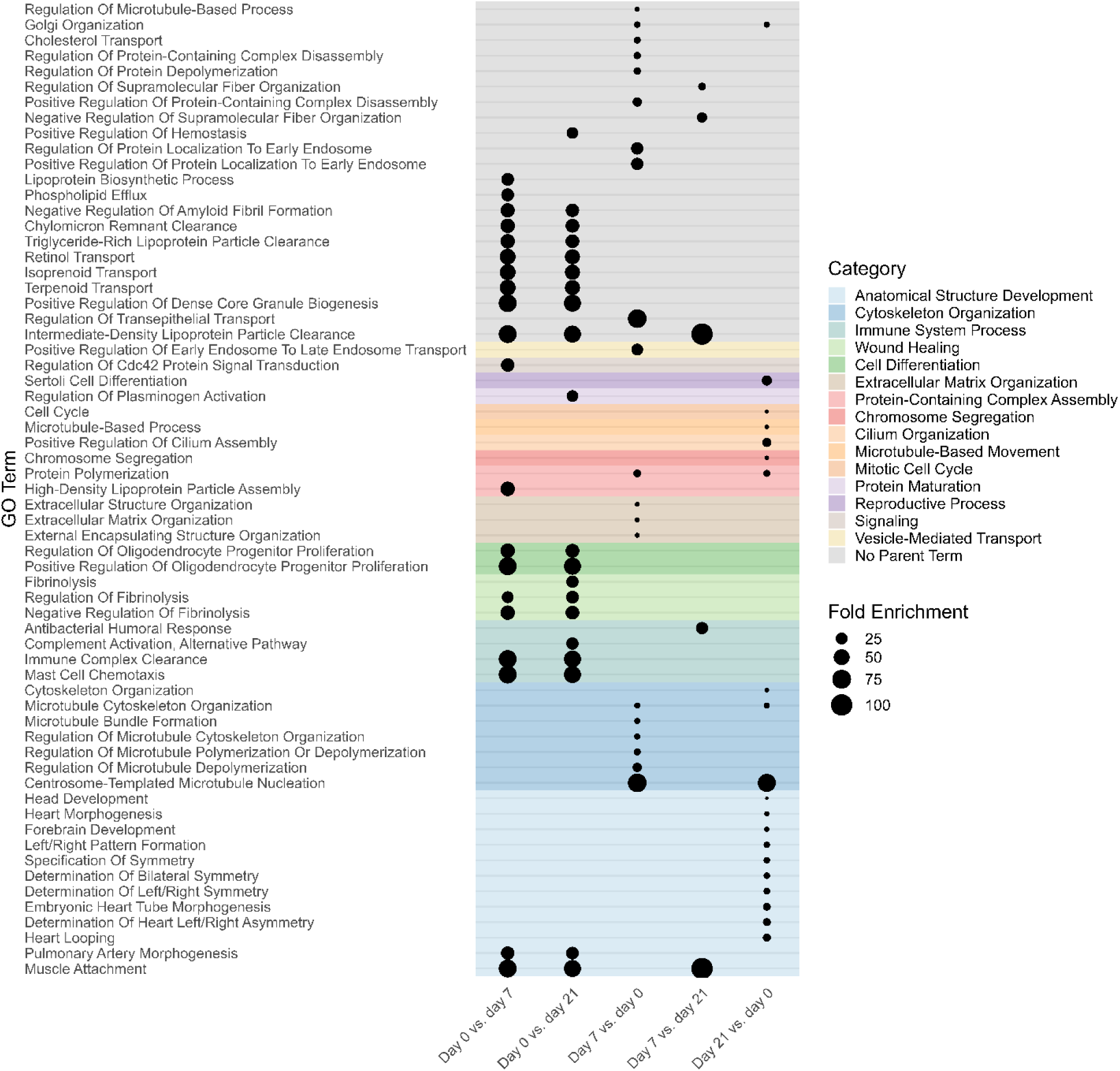
The 20 most significantly enriched gene ontology (GO) biological processes identified in sow colostrum and milk within each pairwise time-point comparison. Dot size represents fold enrichment, while colours indicate functional categories of the GO terms.

A significant but weak correlation was observed between the microbiome and proteome of the colostrum and milk samples (Procrustes correlation = 0.29; *P* = 0.029). Correlations between individual species and proteins were also calculated. The strongest associations included *Rothia nasisuis* with crystallin beta-gamma domain containing 3 (Spearman’s *r* = 0.87) and afamin (*r* = - 0.85) and *Rothia nasimurium* with SIN3 transcription regulator family member A (*r* = 0.86) and alpha-lactalbumin (r = 0.85) (Fig. 5; Table S11). Other notable positive correlations were observed between mucin-1 and *Moraxella* sp024137865, *R*. *nasimurium*, *S*. *suis* (V), and UBA9076 sp022772735. The relative abundance of the swine pathogen *Glaesserella parasuis* correlated with several proteins, including polynucleotide adenylyltransferase (*r* = 0.83), apolipoprotein E (APOE) (*r* = -0.82), granulin (*r* = -0.81), tenascin C (*r* = 0.81), and afamin (*r* = -0.80).

**Figure 5.**
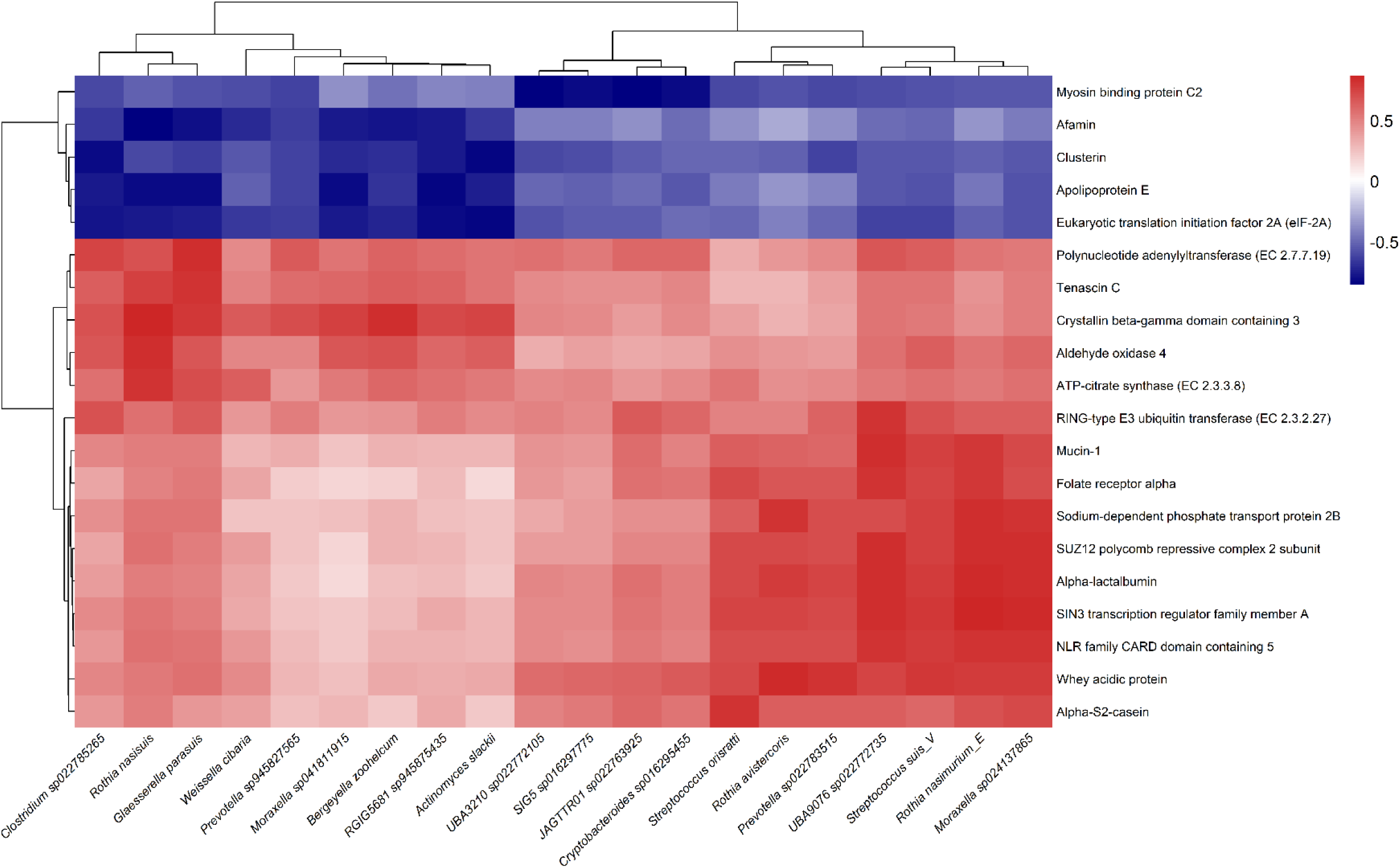
Heatmap of Spearman correlation coefficients between the 20 proteins and 20 bacterial species most frequently represented among the top 500 significant protein–species correlations (FDR < 0.05).

## DISCUSSION

In addition to serving as a primary source of nutrients and immunoglobulins, sow colostrum and milk represent one of the earliest microbial exposures for the piglet gastrointestinal tract. Although previous studies have isolated bacteria from sow colostrum and milk [6] and profiled the colostrum and milk microbiota [5, 10, 11], their reliance on 16S rRNA gene sequencing for microbial identification and characterization is limiting. To address this knowledge gap, we characterized the microbiome of sow colostrum and milk using both culturomics and metagenomic sequencing. In parallel, we analyzed the host-derived proteomes of these samples and investigated potential associations between the milk microbiome and host proteome.

We recovered 132 bacterial isolates representing at least 42 unique species, including 15 putative novel species. The collection included several potentially beneficial bacteria such as *L*. *amylovorus*, *L*. *curvatus, Ligilactobacillus salivarius, L. plantarum,* and *L. reuteri.* Among these, the *L*. *amylovorus* isolates were the most relatively abundant in the colostrum and milk metagenomes and were also relatively abundant in piglet fecal microbiomes during the nursing period, suggesting potential maternal transfer. Certain *L*. *amylovorus* strains have been shown to reduce adhesion of enterotoxigenic *E*. *coli* (ETEC) and mitigate epithelial cell damage *in vitro* [52, 53] and to lower the abundance of ETEC in the ileum of pigs *in vivo* when delivered via feed in challenge experiments [54]. Similarly, *L*. *reuteri*, which was also relatively abundant in colostrum and milk, has been linked through dietary supplementation to increased average daily gain and reduced diarrheal incidence in pigs [55]. As such, it is clear that sow colostrum and milk serve as an early source of beneficial lactobacilli that contribute to piglet health and development.

While sow colostrum and milk contain beneficial bacteria such as the lactobacilli, they may also be a source of pathogenic or opportunistic species, including isolates identified as *A*. *suis*, *S*. *hyicus*, *S*. (*Clostridium*) *perfringens*, and *S*. *suis*. The only publicly available genome of *A*. *suis* beyond our isolate is that of the type strain, originally obtained from the brain of a pig with meningitis [56]. *S*. *hyicus* is the main etiologic agent of exudative epidermitis, also known as “greasy pig disease” [57], *S*. *suis* is a leading cause of meningitis and sepsis [58], and *S*. *perfringens* can cause necrotic enteritis in piglets, depending on toxinotype [59]. However, these species are also part of the commensal microbiome in swine and their detection may therefore reflect normal colonization rather than pathogenic activity. Consistent with this, *S*. *perfringens* was relatively abundant in the piglet gastrointestinal tract (> 0.5%) yet all piglets remained healthy.

Antimicrobial resistance remains a serious challenge in both human and veterinary medicine. Sow colostrum and milk may represent a potential route for the transfer of antimicrobial-resistant bacteria to piglets. Therefore, in the present study we screened the isolates for known ARGs, focusing on those associated with mobile genetic elements such as plasmids. As expected, the most frequently detected ARGs were those corresponding to antimicrobials historically used in swine production, including the aminoglycosides, macrolides, and lincosamides, and tetracyclines. Several ARGs were identified on the same contig as plasmid replicons. Among these was *tet*(K), a tetracycline efflux pump gene co-located with rep7a in a number of *S. epidermidis* and *S. simulans* isolates. The *tet*(K) gene has previously been found with rep7a in multiple *Staphylococcus* species, including *S*. *epidermidis* from pigs and swine farmers [60].

One *L*. *curvatus* isolate (11269D021BM) carried *fexB* and *poxtA* together on a plasmid (rep29) and *tet*(M) on a separate plasmid (rep28). Both *fexB* and *poxtA* confer resistance to florfenicol which is used in swine production to prevent and treat bacterial infections. However, *poxtA* also mediates resistance to the oxazolidinones [61], an antimicrobial class not approved for use in veterinary medicine as they are considered medically important for humans. The oxazolidinones include linezolid, a drug often reserved for treating serious infections such as MRSA and VRE [48, 62]. Recently, we also identified *fexB* and *poxtA* co-located on a plasmid in florfenicol- and linezolid-resistant *Enterococcus avium*, *Enterococcus faecalis*, and *E*. *faecium* isolates recovered from piglets treated with florfenicol [15]. Plasmid-encoded *fexB* and *poxtA* have also been detected in a *L. salivarius* strain isolated from a pig [63]. These findings emphasize the need for careful screening of lactobacilli strains for ARGs during probiotic development.

In addition to the cultured isolates, we assembled MAGs from the colostrum and milk metagenomes. These included species such as *B*. *zoohelcum* and *Sodaliphilus* sp004557565 from the phylum Bacteroidota, as well as *B. mongoliense,* which were not recovered through culture-based methods. This likely reflects limitations in the media and culturing conditions used, as no members of the Bacteroidota were isolated. Although seemingly rare, *B*. *zoohelcum* was recently associated with respiratory disease in pigs [64], while *B. mongoliense* has been found in bovine milk and raw milk cheese with several strains able to degrade milk oligosaccharides [65]. In total, 9 of the 18 high-quality MAGs assembled represented archaeal or bacterial species without a cultured representative. Notably, one MAG (SUG3746) was assigned to the same novel species (*Rothia* sp034179845) as two isolates (32237D007AR and 88186D007BW). Several other MAGs, including *L*. *amylovorus* SUG3751, *S*. *hyovaginalis* SUG3747, and *S. ventriculi* SUG3742, were closely related to at least one isolate, demonstrating the value of MAGs in capturing both cultured and uncultured members of the colostrum and milk microbiome.

The microbial species and functional composition of the colostrum shifted significantly as the milk matured. The transition from colostrum (day 0) to early mature milk (day 7) was marked by changes in the relative abundance of a large number of archaeal and bacterial species, as well as KEGG functional pathways encoded by these microbes. *Rothia* sp034090985 was among those species with the largest relative increase in day 7 and 21 milk compared to colostrum. However, other *Rothia* spp. were relatively abundant in the colostrum, including *Rothia* sp034179845, a placeholder species from which we recovered two isolates. Other abundant *Rothia* spp. were *R*. *nasimurium and Rothia endophytica*. *Rothia* spp. are commonly reported among the abundant bacteria in sow colostrum and milk [66], as well as in the porcine tonsil [67], upper respiratory tract [68], oral cavity [11], and skin [69] microbiomes. They are present in the pig gastrointestinal tract as well, although typically at a lower relative abundance compared with other body sites [70]. Although certain *Rothia* spp., such as *R*. *nasimurium* isolated from pigs, have been previously reported to carry CAZymes that can metabolize milk oligosaccharides and mucins [71], none of the *Rothia* spp. isolate genomes in the current study encoded CAZymes specifically associated with milk oligosaccharide degradation. Milk oligosaccharides are glycans composed of a lactose core linked to monosaccharides such as fucose and N-acetylglucosamine, as well as sialic acids including N-acetylneuraminic acid and, in pigs, N-glycolylneuraminic acid [72]. These oligosaccharides are notable because they are not typically digested by the piglet but are instead metabolized by specific members of the gut microbiome, thereby providing benefits to the host such as selective stimulation of beneficial bacteria, production of short-chain fatty acids (SCFAs), and modulation of the immune system. Due to structural similarities, many of the CAZymes in the milk oligosaccharide-associated GH families may also be involved in the degradation of other host glycans such as mucins [73]. Mucins are high-molecular-weight glycoproteins that constitute the major structural components of mucus lining the epithelial surfaces of mammals; consequently, the ability to metabolize mucins can enhance bacterial colonization of the host [74].

Among the bacterial isolates, the *S*. *perfringens* genomes encoded the greatest number of CAZyme families linked to milk oligosaccharide metabolism. *In vitro*, *S*. *perfringens* has been shown to metabolize several human milk oligosaccharides, producing SCFAs such as butyrate and propionate [75]. Although none of the three *S*. *perfringens* isolate genomes were relatively abundant in the colostrum or milk, they were relatively abundant in the piglet fecal metagenomes suggesting that the presence of these CAZymes, if expressed, may provide an early colonization advantage through the degradation of dietary milk oligosaccharides or host mucins. Lactose is the major carbohydrate in colostrum and milk, and tends to increase in concentration over the lactation period [76]. Given this, it is not surprising that nearly all isolate genomes encoded at least one GH family associated with β-galactosidase activity, enabling hydrolysis of lactose into glucose and galactose.

We also characterized the milk proteome and examined its association with the microbiome to better understand potential host-microbe interactions. In agreement with previous studies on sow colostrum and milk, proteins such as APOE, APOA4, APOH, ITIH2, PAPLN, TIMP2, and TF were associated with colostrum and LALBA, LPO, and MUC1 with mature milk [77, 78]. These proteins reflect the transition from colostral proteins involved in nutrient and lipid transport and immunity to mature milk proteins associated with antimicrobial activity and lactose synthesis. Also among the most strongly colostrum-associated proteins was a colostrum trypsin inhibitor-like protein, the gene for which has been reported to be expressed over sevenfold higher in sow colostrum compared to mature milk [79]. This protein likely helps protect colostral immunoglobulins from degradation by trypsin secreted by the piglet small intestine [79]. The concentration of mucin-1, a transmembrane mucin, was strongly correlated with the relative abundance of *R. nasimurium*. *Rothia* spp. typically produce enzymes capable of degrading mucins [71]; however, as mentioned earlier, the *Rothia* spp. isolated in the present study lacked most CAZymes required for host glycan metabolism.

*Glaesserella parasuis*, a swine pathogen, was positively correlated with tenascin C. This extracellular matrix glycoprotein can bind to many different ligands, including toll-like receptor 4 (TLR4), initiating the release of pro-inflammatory cytokines. *G*. *parasuis* was also negatively correlated with afamin, a glycoprotein that has been found to be decreased in the serum of patients with certain infections or diseases [80]. Similarly, the relative abundance of *G*. *parasuis* was also negatively correlated with APOE, a lipid-binding protein. Interestingly, APOE has been shown to have antibacterial activity against certain gram-negative bacteria *in vitro* and *in vivo* in mice [81]. These correlations suggest a potential association between *G*. *parasuis* abundance and host inflammatory and defence proteins, consistent with a possible localized host response that may influence bacterial persistence in milk.

The exact source of bacteria present in mammalian colostrum and milk remains unclear. However, oro/entero-mammary translocation from the gastrointestinal tract or oral cavity to the mammary gland has been proposed as a plausible explanation for at least some of the species detected [8, 82]. Despite careful cleaning and disinfection of the teat area prior to sample collection, certain bacterial species identified in this study likely originated from the skin surrounding the teat, either as part of the skin microbiome or through fecal contamination. Regardless of their origin, these bacteria represent microbes that piglets naturally ingest while nursing. As such, these early exposures likely play an important role in shaping the development of the neonatal gut microbiome and influencing immune system maturation.

## CONCLUSION

Using both culturomics and metagenomic sequencing, sow milk and colostrum were found to contain a diverse microbiome that changes during lactation. Although the role of many of these bacterial species remains unclear, some, such as the lactobacilli, likely contribute to piglet health during the nursing period. Given the number of ARGs identified in the bacterial isolates, particularly those associated with plasmids, sow colostrum and milk may represent an early source of ARGs and antimicrobial-resistant bacteria for piglets. The colostrum and milk proteomes showed corresponding shifts in immune- and metabolism-related proteins across lactation, and their correlations with specific microbial taxa suggest potential host-microbe interactions during lactation. This study provides a foundation for future work investigating how colostrum and milk microbiomes influence early-life microbial colonization, and may inform strategies to enhance piglet health through microbiome manipulation.

### Author contributions

D.B.H. designed the research. A.K., D.B.H., and K.E.G. carried out bioinformatics and statistical analyses. D.B.H., K.E.G., and P.M. evaluated proteomic data. D.B.H. wrote the manuscript. A.K., K.E.G., and P.M. reviewed and edited the manuscript. All authors read and approved the final manuscript.

## Acknowledgements

The authors thank the Lacombe Research and Development Centre animal care staff for their excellent care of the pigs and assistance with sampling. We are also very grateful to Cara Service for help with sample collection and the initial culturing of bacteria from the colostrum and milk samples, and to TingTing Liu for performing the metagenomic DNA extractions. We also wish to acknowledge Daniel Llewellyn and Zebedi Odongo for their contributions during the initial analysis of the milk proteome.

## Conflicts of interest

The authors declare that there are no conflicts of interest.

## Funding information

Funding for this project was provided by Agriculture and Agri-Food Canada’s A-Base program.

## Ethical approval

Animals were cared for in agreement with the Canadian Council for Animal Care (2009) guidelines. The Lacombe Research and Development Centre Animal Care Committee reviewed and approved all procedures and protocols involving animals.

## Data availability Data summary

The isolate genomes and metagenome-assembled genomes are publicly available in the National Center for Biotechnology Information’s (NCBI) sequence read archive and genome databases under BioProject PRJNA1134706. Piglet fecal and sow colostrum and milk metagenomic sequences are available under BioProject PRJNA779404.

## Notes

### Competing Interest Statement

The authors have declared no competing interest.

